# Dissection of intestines from larval zebrafish for molecular analysis

**DOI:** 10.1101/493536

**Authors:** Bilge San, Marco Aben, Gert Flik, Leonie M. Kamminga

## Abstract

Epigenetic data obtained from whole zebrafish embryos or larvae may mask or dilute organ-specific information. Fluorescence activated cell sorting can diverge cells from their native state, and cryosections often yield insufficient material for molecular analysis. Here, we present a reproducible method for larval intestinal isolation at 5, 7, and 9 days post-fertilization, using the intestine-specific transgene *tgBAC(cldn15la:GFP)*. With tweezers, the intestine can be pulled out of the abdomen in one smooth motion. Upon removal of adhering tissues, intestines can be directly used for analyses. Each dissection takes 3-6 minutes per fish. We demonstrate that 10 and 25 dissected intestines yield enough material for RNA-sequencing and ChIP-sequencing, respectively. This method results in high quality, live material, suitable for many downstream applications.

**METHOD SUMMARY:** We present a reproducible method for zebrafish larval intestinal isolation which results in high quality, live material. With tweezers, the intestine can be pulled out of the abdomen and after removal of adhering tissues, intestines can be directly used for analyses. We demonstrate that 10 and 25 dissected intestines yield enough material for RNA-sequencing and ChIP-sequencing, respectively.

## INTRODUCTION

Genetic and epigenetic studies on zebrafish embryos and larvae require different, stage-dependent approaches. Whole embryo lysates are commonly used for studies on gene expression and epigenetics during early embryonic development [1–3]. However, as tissues and organs are specified, information originating from a defined tissue may mask another and signals may ‘dilute’. To eliminate noise and increase reliability, isolation of specific tissues or cells of an organ becomes mandatory.

To obtain organ-specific information for whole genome (DNA) or transcriptome (RNA) analysis, tissues can be dispersed and the cells sorted by fluorescence activated cell sorting (FACS) [4]. FACS enables the collection of specifically labeled living (single) cell populations out of a whole tissue or organism. It is a broadly applied method, for instance for blood cell subtyping [5]. However, cell surface markers might behave differently in single cell suspensions and might be cleaved by proteases (*e.g.* Trypsin) [6]. In zebrafish, unlike mammals, there is limited availability for commercial antibodies for cell surface markers, therefore, FACS is commonly used with transgenic lines which express tissue-specific fluorescent proteins. Cells obtained by FACS can then be pooled for DNA (chromatin), RNA, protein extraction, or separated for single cell studies [7]. Long preparation times, however, decrease cellular yield [8] and lead to anoikis (*i.e.* apoptosis caused by absence of cellular contacts) [9]. Importantly, single cells from dissociated tissues may undergo transcriptional changes, including immediate early response gene activation (e.g. *fos, jun, hsp* gene variants) [10] and further alterations in cell signaling pathways [11]. These alterations in dissociated cells can also cause dedifferentiation [12].

As an alternative to FACS, transcriptome of serial (cryo)sections of whole zebrafish embryos can be sequenced to generate a gene expression map (*e.g.* by Tomo-seq [13]). However, (cryo)sectioning may also cause alterations from native cellular conditions. To assess gene expression in only a subset of cells, cells can be extracted from tissue sections by carbon dioxide laser capture microdissection [14]. These methods are limited to RNA- and DNA-sequencing; a (part of a) single embryo or larva is currently insufficient for chromatin immunoprecipitation with commercially available antibodies, independent of the stage of (early) development [15].

Dissection of organs is a common procedure in studies on adult zebrafish [16], while embryonic and larval dissection studies are uncommon or not well-described. The embryonic heart is the most commonly dissected organ due to its peripheral position in the body, and the broad research interest in its regenerative capacity [17-21]. The zebrafish pronephros, precursor of kidney tissue among others, is another organ which has been dissected at 5 days post-fertilization (dpf) to study gene expression by real-time quantitative PCR [22]. Zebrafish intestine is of great interest due to its rapid development, renewal potential, and its function in supplying nutrients to the larvae after yolk depletion. A number of laboratories have documented intestinal dissections, however, either the dissection method used has not been explained in detail, or the subsequent technical analysis did not require a pure intestine [23-26]. Therefore, we investigated the feasibility of intestinal dissections in zebrafish larvae.

Zebrafish intestinal development begins with the appearance of an array of endodermal epithelial cells along the ventral midline of the embryo between 1-2 dpf [27]. This array of cells gradually forms a single, continuous lumen by the hollowing and subsequent fusion of several small lumina between 2-3 dpf [28]. During intestinal lumen formation, the liver and pancreas ‘Anlagen’ differentiate at the junction between the esophagus and the intestine and go through extensive remodeling and proliferation [27]. Zebrafish is a stomachless species, and its intestine is clearly separated into three parts: intestinal bulb, mid-intestine, and posterior intestine. By 5 dpf, the intestine becomes functional with the opening of the mouth and anus, when most yolk is absorbed and the larva starts feeding exogenously. To understand the regulatory processes in such a rapidly developing organ, the analysis of different time-points becomes necessary. Throughout the second week of development, different epithelial cell subtypes, namely enterocytes, goblet cells, enteroendocrine cells, and specialized antigen presenting (NaPi+) enterocytes (in order of abundance) continue to differentiate [29,30]. With the growth of the intestine, this anterior loop folds into a sigmoid shape by adulthood [31]. Although the zebrafish intestinal lining (an epithelium very rich in enterocytes) is very similar in structure to that of mammals, it has ridges instead of villi. Proliferation, like in mammals in the crypts, predominantly occurs at the base of these ridges [30].

We present a rapid and reproducible method of dissection of larval zebrafish intestine with the aid of the intestine-specific transgenic line *tgBAC(cldn15la:GFP)*, which expresses the GFP-tagged protein ‘claudin 15-like a’, an integral protein in the tight junctions of the intestinal epithelium [28]. We show that this technique is compatible with methods such as RNA- and ChIP-sequencing, and surpasses the efficiency of FACS of intestinal cells in the larval stages of this transgenic line.

## MATERIALS AND METHODS

### Zebrafish strains and husbandry

Transgenic lines *tgBAC(cldn15la:GFP*) [28] and *tg(gut:GFP)* [32] were used for GFP expression in the intestine. Embryos were raised in E3 embryo medium at 28.5°C as described in detail elsewhere [33]. GFP expression was checked at 3-5 dpf under light anesthesia in 2-phenoxyethanol (0.05% v/v). Larvae were fed twice daily with dry feed (Gemma Micro 75, Skretting), rotifers, and artemia according to guidelines [33,34]. All experiments described are in accordance with institutional animal welfare guidelines, policies, and laws, and were approved after ethical testing by Central Committee for Animal Experimentation (CCD, approval number AVD1030020184668) of the Netherlands.

### Fluorescence Activated Cell Sorting

Two hundred 5, 7 or 9 dpf larvae in the *tg(gut:GFP)* or *tgBAC(cldn15la:GFP)* background were anesthetized, divided into 1.5 ml microcentrifuge tubes (20 larvae per tube), and washed with PBS. The larvae were dissociated in 0.25% w/v Trypsin (Sigma), 1 mM EDTA in PBS at 28.5°C for 50-70 minutes. After trypsinization was stopped with 1 mM CaCl_2_ and 100 µl 100% FBS, the suspension was passed through a FACS filter (BD, 70 µm) and clarified with Dnase I (100 µg/ml). Finally, the suspension was washed two times in PBS/1 mM EDTA solution and stained with 7-Aminoactinomycin D (7-AAD, Thermo Fisher) for cell viability. GFP-positive cells were sorted by BD FACS-Aria into TRIzol (Thermo Fisher).

### Dissection

A Petri dish lid was positioned under a fluorescence stereo microscope (Leica MZ FLIII) as a working surface. During all steps, light microscopy and fluorescent microscopy were combined. Up to 4 *tgBAC(cldn15la:GFP)* larvae of 5, 7, or 9 dpf were placed in 2-phenoxyethanol (0.05% v/v) for anesthesia and processed within 30 minutes following loss of startle response. One larva was transferred under the microscope in 3-4 ml anesthesia medium, facing the dominant hand of the researcher. With one clean watchmaker’s tweezer, the larva was pierced rostrally to the intestine behind the branchial arches and the intestine was clamped. At the same time, the fish was stabilized by pinching the swim bladder with another watchmaker’s tweezer (Figure 1A and 1A’). Next, in one movement, the intestinal tract was carefully and slowly pulled out in the direction of the head; *i.e.* held by the distal segment and pulled towards the head of the fish (Figure 1B and 1B’). The intestinal tract was bisected at the transition between the esophagus and the intestine, and the carcass was discarded or lysed for genotyping (Figure 1C and 1C’). The intestine was cleaned up under the microscope in fresh medium (Figure 1D and 1D’). In 5 dpf larvae, yolk remnants were removed. At all time points, the swim bladder was pinched off with tweezers. Then, with tweezers and a microsurgical blade, liver and pancreas connections were cut off to free the intestine. Remnants of muscles were peeled off with tweezers. The intestine was double-checked for remaining adhering tissues (*e.g.* the autofluorescent gallbladder). The pure intestine was washed in clean system water, and then transferred into a 1.5 ml microcentrifuge tube with a Pasteur pipette. The working surface was frequently refreshed. After dissection, a maximum of 15 intestines were pooled on ice to prevent tissue damage before processing.

**Figure 1.**
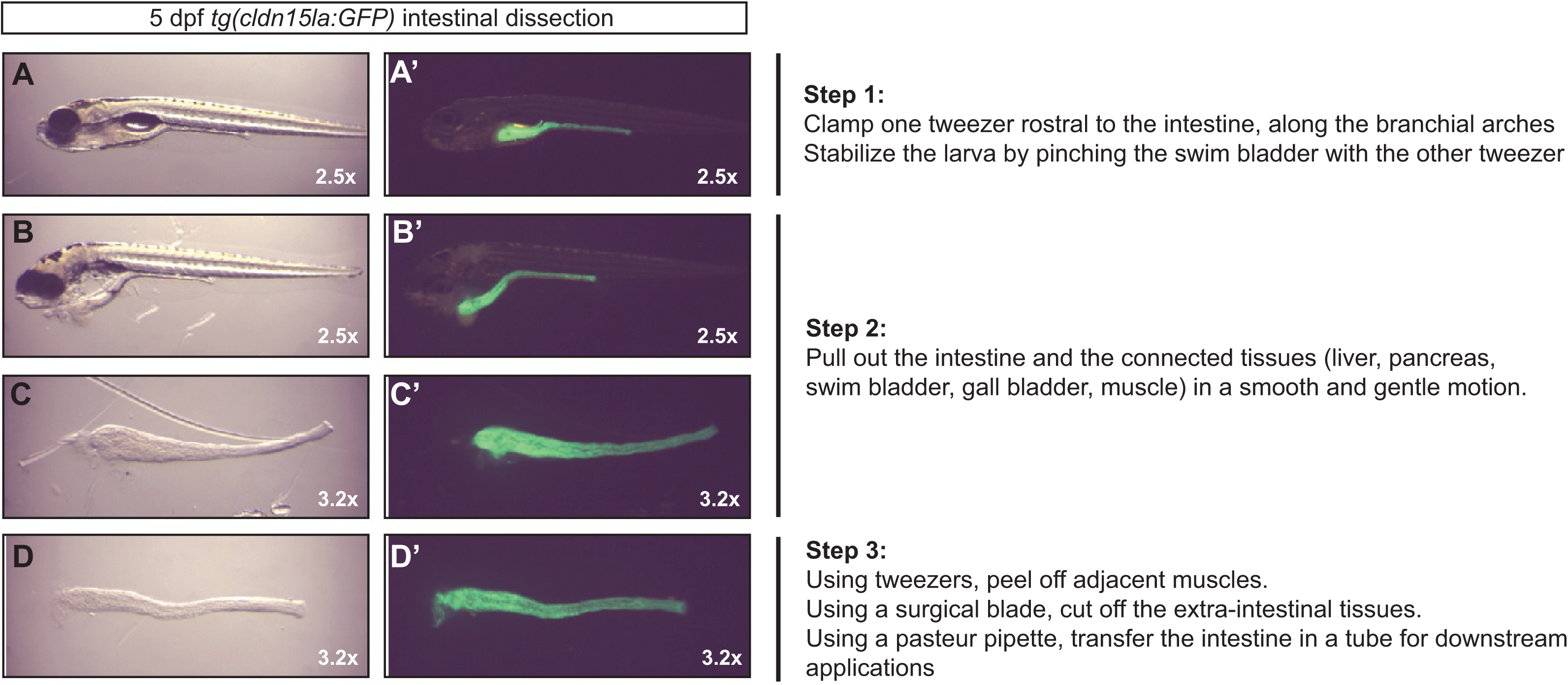
FACS sorting in tg(gut:GFP) and tg(cldn15la:GFP). Single cell suspensions were prepared from 5 dpf whole zebrafish larvae in the *tg(gut:GFP)* **(A)** and *tg(cldn15la:GFP)* **(B)** backgrounds. The GFP-positive populations (**A, B,** purple dots) were gated according to the GFP-negative population (**A, B,** black dots) and sorted into TRIzol for RNA extraction. **C.** FACS on whole *tg(gut:GFP)* and *tg(cldn15la:GFP)* larval suspensions at 5 dpf yielded 0.1% and 1.9% GFP-positive cells, respectively. Unexpectedly, FACS on dissected intestines from *tg(cldn15la:GFP)* yielded as low as 8.9% GFP-positive cells.

### RNA isolation

Microcentrifuge tubes (1.5 ml) were filled with 100 or 500 µl TRIzol for 1 or 10 intestines, respectively. Cells sorted by FACS were collected into 500 µl TRIzol. Dissected intestines were transferred with a Pasteur pipette onto the lid of the tubes with a minimum volume of system water, and promptly shaken in TRIzol for lysis. RNA was isolated as described elsewhere [35]. After phase separation, in-column DNase I treatment was performed (ZYMO Quick-RNA MicroPrep). Total RNA yield was measured by fluorometric quantification (Qubit).

### Chromatin immunoprecipitation

To prevent adsorption of dissected intestines to the tubes, microcentrifuge tubes (1.5 ml) were coated with 5% (w/v) BSA (Sigma) solution [36] for 15 minutes and dried. Thirty dissected intestines were cross-linked in 1% paraformaldehyde (Electron Microscopy Sciences) for 15 minutes. The reaction was quenched with 125 mM glycine for 5 minutes, and washed 3 times with PBS. The intestines were lysed (20 mM Tris-Cl pH 7.5, 70 mM KCl, 1 mM EDTA, 10% glycerol, 0.125% NP40, protease inhibitor cocktail [Roche]) and sonicated (6 cycles of 30 seconds, Bioraptor^®^ Pico) to extract ∼200 bp chromatin fragments. The chromatin was bound to protein A/G beads (Invitrogen, 1003D), incubated with anti-H3K4me3 (Millipore, 2 µg) or anti-H3K27me3 (Millipore, 2 µg) antibodies overnight, then eluted off the beads. Input DNA concentration (1:6 fraction of total) and ChIP yield (5:6 fraction) was measured by fluorometric quantification (Qubit).

## RESULTS AND DISCUSSION

In zebrafish, the intestine, which is the sole deliverer of energy (feed) to the animal, develops very quickly and throughout early larval stages. The organization of the intestine at that moment is comparable to the adult situation. Zebrafish intestinal anatomy is functionally comparable to the anatomy of most higher vertebrates [27]. Zebrafish intestinal epithelium is organized in ridges with somewhat larger dimensions compared to mammalian villus-crypts [30]. Proliferative regulation is similar, *e.g.* with a crucial role for Wnt signaling, which appears conserved from zebrafish to mammals [37,38]. Therefore, the zebrafish intestine is an attractive translational model to study human diseases [39] and for fundamental research on (epi)genetic regulation [1]. For these reasons, we aimed to isolate the intestine from larval stages. This study combines zebrafish developmental physiology with molecular biology and demonstrates a highly feasible technique to dissect the intestinal tract of zebrafish larvae in the *tgBAC(cldn15la:GFP)* transgenic background. It further presents yield of RNA extraction and chromatin immunoprecipitation from intestines, and shows that the total RNA yield from dissected intestine surpasses that of FACS-samples by at least 4-fold in our hands. Here we will discuss the rationale behind the development, advantages, and disadvantages of this method.

### Fluorescence Activated Cell Sorting (FACS)

To study zebrafish intestinal (epi)genetics during the first days of larval development, we used two previously described transgenic lines, *tg(gut:GFP)* [32] and *tgBAC(cldn15la:GFP)* [28]. The rationale was that from these transgenes cell suspensions could be made, from which intestinal cells could be isolated by FACS for further molecular analyses. The isolation of cells from both of these lines prior to FACS analysis presented challenges. Preparation of isolated cell suspensions from whole larvae took over 2 hours, during which cell viability decreased to 60%. GFP-positive and negative cells did not present a distinct boundary to set reliable gates for sorting. The percentage of GFP-positive cells obtained from 5 dpf larvae was as low as 0.1% for *tg(gut:GFP)* (Figure 2A), and 1.9% for *tgBAC(cldn15la:GFP)* (Figure 2B). As the *tg(gut:GFP)* line also expresses the construct in the liver and pancreas and gave such low FACS yield, we decided the continue our research with the *tgBAC(cldn15la:GFP)* line.

**Figure 2.**
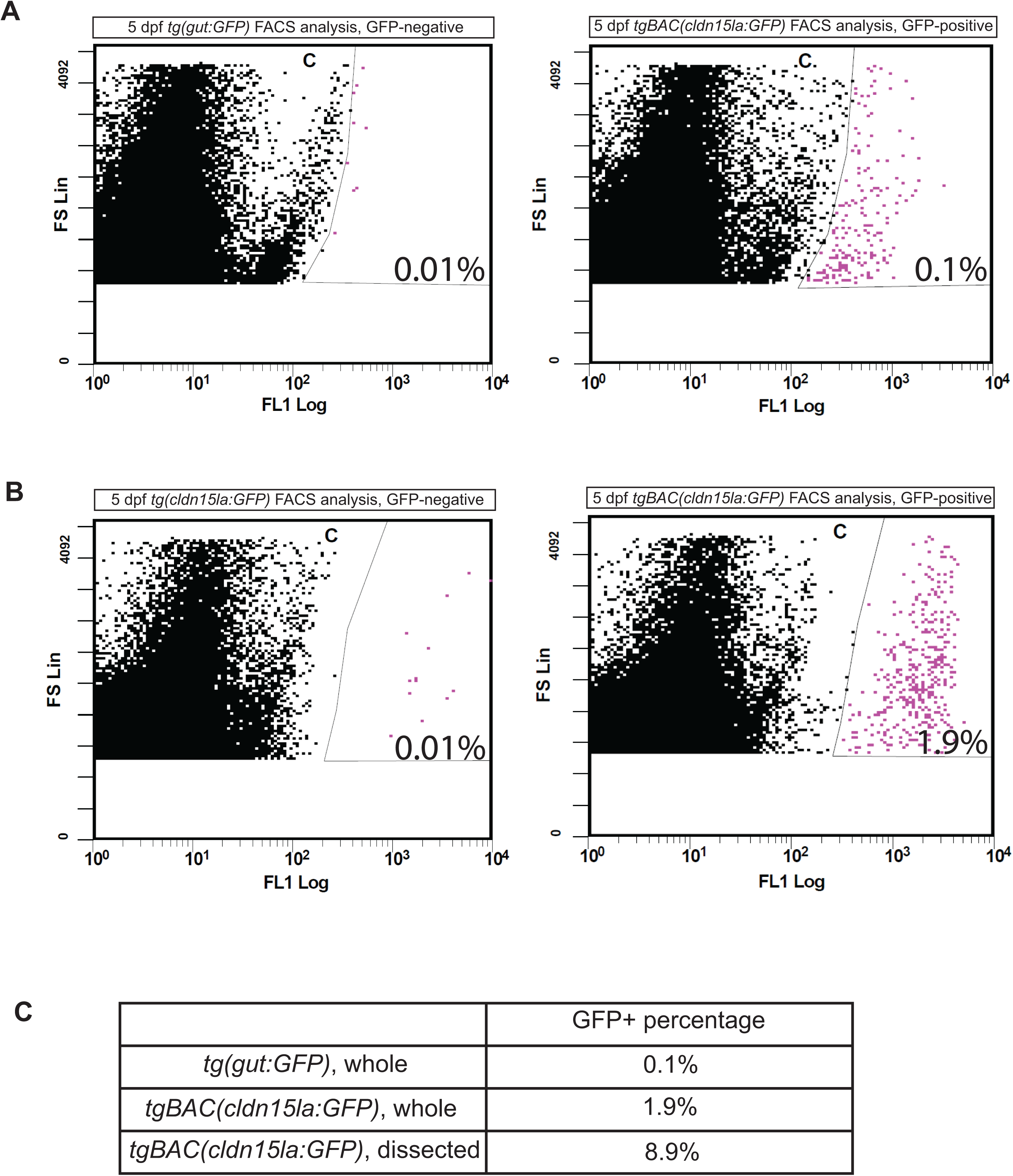
Dissection of the larval intestine. Overview of the major steps during the dissection of a larval intestine; 5 dpf is shown as an example. The same steps are taken for 7 and 9 dpf. Left and right panels are the same field of view under light microscopy (A, B, C, D) and fluorescent microscopy (A’, B’, C’, D’), respectively. **A, A’.** With the help of tweezers, the intestine was stabilized. **B, B’.** The intestine was slid out of the body in the direction of the head. **C, C’.** The extra-intestinal tissues were cleaned off by peeling or cropping by microsurgical blades. **D, D’.** The intestine was carefully checked for GFP purity and promptly transferred into a microcentrifuge tube.

Cell dissociation for FACS is a rigorous process for cells due to the stress of enzymatic digestion and trituration. Hard and soft tissues require different durations to dissociate, and the timing of complete dissociation changes according to the age of embryos/larvae. During cell dissociation, some cell death occurs and the viscous texture of genomic DNA in solution may encumber pipetting. As mentioned before, changes might occur in transcription, cell signaling pathways, and the differentiation status of the cells [10-12].

### Intestinal dissection

Next, we investigated whether segmenting the larvae into smaller parts prior to single cell preparations would increase the FACS yield and reduce noise from auto-fluorescence. For this inquiry, we attempted dissections on larval intestines in the *tgBAC(cldn15la:GFP)* background. Remarkably, these dissections were very consistent and time efficient. Moreover, the reduction of material processed per zebrafish greatly reduced the trypsinization time as well; approximately by half. Nonetheless, the FACS yield of this semi-pure population of GFP-positive cells was only 8.9% at 9 dpf (Figure 2C, Supplementary Figure 1). Due to this (unexpected) low yield, we concluded that *tgBAC(cldn15la:GFP)* is an unsuitable model for FACS. However, it could serve as a great tool for obtaining intestine-specific cells through dissections.

Dissection of the intestine resulted in minimal tissue damage due to the short processing time (3-6 minutes per fish) required. To prevent loss of sample quality and to obtain intestines in a comparable developmental stage per batch, we limited handling time to 60 minutes, or 15 larvae, before lysis or fixation. During dissection, the integrity of the intestine was visualized in real-time by microscopy. This allows possible (mutant) intestinal phenotypes or technical errors in dissection to be observed under the microscope. Moreover, the carcass from each larva can be genotyped, and any single unsuitable sample can be discarded before intestinal pooling.

During dissection optimizations, sliding the intestine out of the body in rostrocaudal direction proved to be the simplest, fastest, and the most reproducible method. Each intestine took 3 to 6 minutes to collect, mainly depending on the age of the larva (and the experience of the researcher); older larvae could be dissected in shorter time. Once the intestines were out of the larvae, non-intestinal tissues were removed under fluorescent and light microscopy. This reproducible purification of intestinal epithelium proved to be an excellent start for molecular analysis, as opposed to a mix of tissues in whole larvae.

During the development of the technique, several aspects of the protocol needed to be considered for optimal results. To manipulate larvae easily under the microscope and to minimize light refraction, the system water on the working surface (petri dish lid/cover) was set between 3-4 ml. To prevent the adherence of intestines to plastic, glass Pasteur pipettes were used for transfer of intestines, and BSA coating was chosen over the rather costly and inefficient use of dichlorodimethylsilane coating or low binding microentrifuge tubes. Bovine serum albumin (BSA) is a protein commonly used for blocking Western blot membranes and ChIP beads, but also for coating laboratory equipment against adherence of materials [40].

Intestinal dissections can be considered as a difficult process prone to errors and variation. Between 5-9 dpf, the length of the larvae is between 3.9-4.5 mm, which requires the use of a microscope and watchmaker’s tweezers. For the tissue to stay intact and unchanged from the start of the dissection to tissue lysis/fixation, the researcher needs to act fast and gentle at the same time. Individual variation in physiology [41] also needs to be considered for analysis of multiple fish.

In addition to individual differences, time point differences also unavoidably vary the dissection procedure. Because the intestinal tissue is still soft and elastic at 5 dpf [34], it is more likely to tear between the intestinal bulb and mid-intestine during the removal of the intestine from the body. The muscles are the most challenging extra-intestinal tissue to remove at 5 dpf due to the fragility of the intestine and the thinness of the muscle lining. By 7 dpf, the intestine has hardened enough such that muscle, liver, and pancreas can be swiftly peeled/cut off of the intestine with the help of tweezers and microsurgical blades.

Further, the procedure is potentially prone to contamination with liver, pancreas, muscle, and gallbladder. At 5 dpf, yolk contamination is also an additional risk. Therefore, at least 6 biological replicates should be used if the extracted RNA will be used for sequencing [42]. For ChIP-sequencing, more than two replicates are recommended to minimize errors in bioinformatics analysis [43]. Another aspect of sample variation is the presence of four different epithelial cell types and three different morphological segments within the intestine [29,30], with different functional properties [31]. It is a coherent presumption that these different functions start developing before or during larval stages.

### RNA isolation and chromatin immunoprecipitation

We used dissected intestines for RNA isolation and chromatin immunoprecipitation. At all time points, total RNA obtained by single intestines (average: 28.4 ng) and pools of 10 intestines (average: 343 ng) was proportional to the number of larvae (Table 1). Surprisingly, FACS on intestinal cells at 9 dpf from 20 larvae yielded a proportionally >4-fold lower amount of total RNA than dissected intestines (Table 1). The amount of RNA from dissected pooled intestines suffices as input for RNA-sequencing (Ribo-Zero rRNA Removal Kit, Illumina) for all developmental time points used.

**Table 1.** Total RNA yield. Total RNA was isolated from single or a pool of 10 dissected intestines at 5, 7, and 9 dpf in triplicates, and the yield was quantified fluorometrically (Qubit). The average yield is shown in nanograms. FACS-sorted intestinal cells from 20 whole *tg(cldn15la:GFP)* larvae at 10 dpf yielded 4-fold less total RNA.

|  | 5 dpf | 7 dpf | 9 dpf |
| --- | --- | --- | --- |
| Single intestines | 24.4 | 30.8 | 29.9 |
| 10 intestines | 364.5 | 348 | 316.5 |
| FACS (20 larvae) | N/A | N/A | 150 |

To immunoprecipitate intestinal chromatin, we collected intestines in a BSA-coated microcentrifuge tube. Immunoprecipitation of chromatin from 25 pooled intestines with anti-H3K4me3 and anti-H3K27me3 antibodies yielded sufficient starting material for Illumina sequencing preparation; on average 6.3 ng and 12.8 ng chromatin was immunoprecipitated with anti-H3K4me3 and anti-H3K27me3, respectively (Table 2). We have used total RNA and chromatin from pooled zebrafish larval intestines to analyze the wild type intestinal transcriptome and the presence of H3K4me3 and H3K27me3 chromatin marks on gene promoters at 5, 7, and 9 dpf [44]. To detect individual variation, single intestines should be processed with available low input protocols [45-47]. However, low-input ChIP yields still predominantly depend on the antibody efficiency, and the methods are costly for many [48].

**Table 2.**
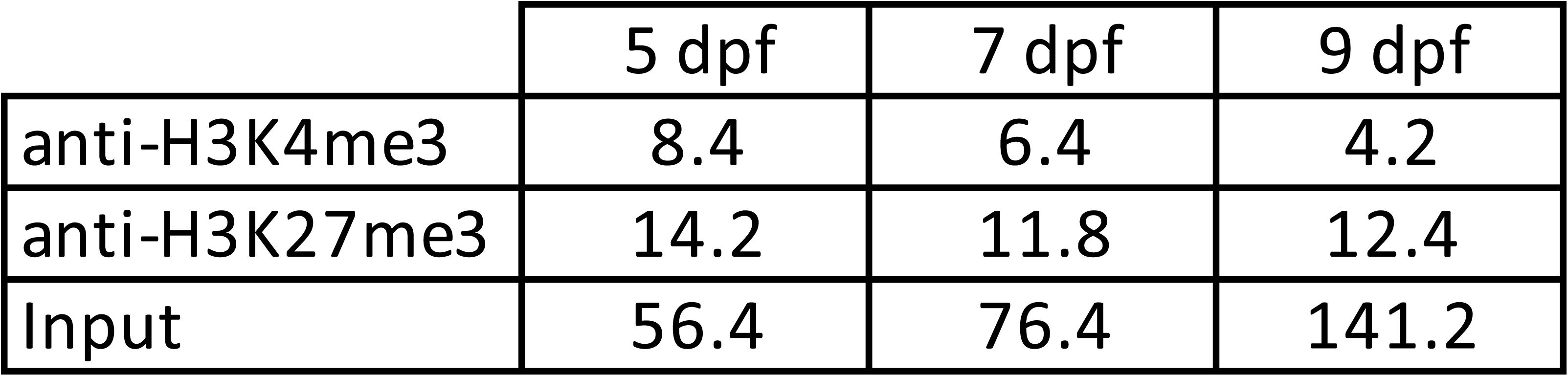
Chromatin immunoprecipitation yield. After chromatin extraction from 30 intestines at 5, 7, or 9 dpf, samples were sonicated to obtain ∼200 bp fragments, and one sixth of the DNA (∼5 intestines) was separated to measure DNA input. The rest of the sample (five sixth, ∼25 intestines) was subjected to anti-H3K4me3 or anti-H3K27me3 immunoprecipitation in replicates, eluted off, and the yield was quantified fluorometrically (Qubit). The average ChIP yield is shown in nanograms.

|  | 5 dpf | 7 dpf | 9 dpf |
| --- | --- | --- | --- |
| anti-H3K4me3 | 8.4 | 6.4 | 4.2 |
| anti-H3K27me3 | 14.2 | 11.8 | 12.4 |
| Input | 56.4 | 76.4 | 141.2 |

Although dissected intestines are a far better model than whole larvae for molecular and biochemical analysis of the intestine, we recommend additional validation experiments such as staining of individual mRNA or proteins to assess their localization. Additionally, dissected intestines can be used for protein isolation and subsequent proteomics. In summary, intestinal dissection serves as an excellent tool to compare differences in this rapidly developing organ between larval stages, and between wild types and mutants.

## Supporting information

## AUTHOR CONTRIBUTIONS

B.S. conceived and designed the methodology and performed the experiments, analyzed the data, and prepared figures. M.A. performed experiments. L.M.K. conceived and supervised the study, acquired funding, was responsible for project management. B.S., G.F., and L.M.K. wrote the manuscript. All authors reviewed the manuscript.

## ACKNOWLEDGEMENTS

The authors would like to thank Tom Spanings and Antoon van der Horst of Radboud University for zebrafish husbandry, Rob Woestenenk of the Radboud University Medical Center for FACS assistance, Cornelia Veelken of the Radboud Institute for Molecular Life Sciences for sharing the FACS protocol, Karolina Andralojc of the Radboud University for discussions on BSA-coating, Silvia Boj of the Hubrecht Institute for the *tg(gut:GFP)* line, Ashley Alvers Lento and Michel Bagnat of Duke University for providing the *tgBAC(cldn15la:GFP)* line.

## FIGURE LEGENDS

**Supplemental Figure 1. *FACS sorting in dissected tg(cldn15la:GFP) intestines.*** Intestines of 10 dpf were crudely dissected from *tg(cldn15la:GFP)* larvae and single cell suspensions were prepared. GFP-negative population (left panel, red dots) was used to gate the GFP-positive population (right panel, red dots). GFP-positive population was calculated as 8.9%.

