## Supplementary figures and images for "Dissection of intestines from larval zebrafish for molecular analysis"

### Supplementary file 1

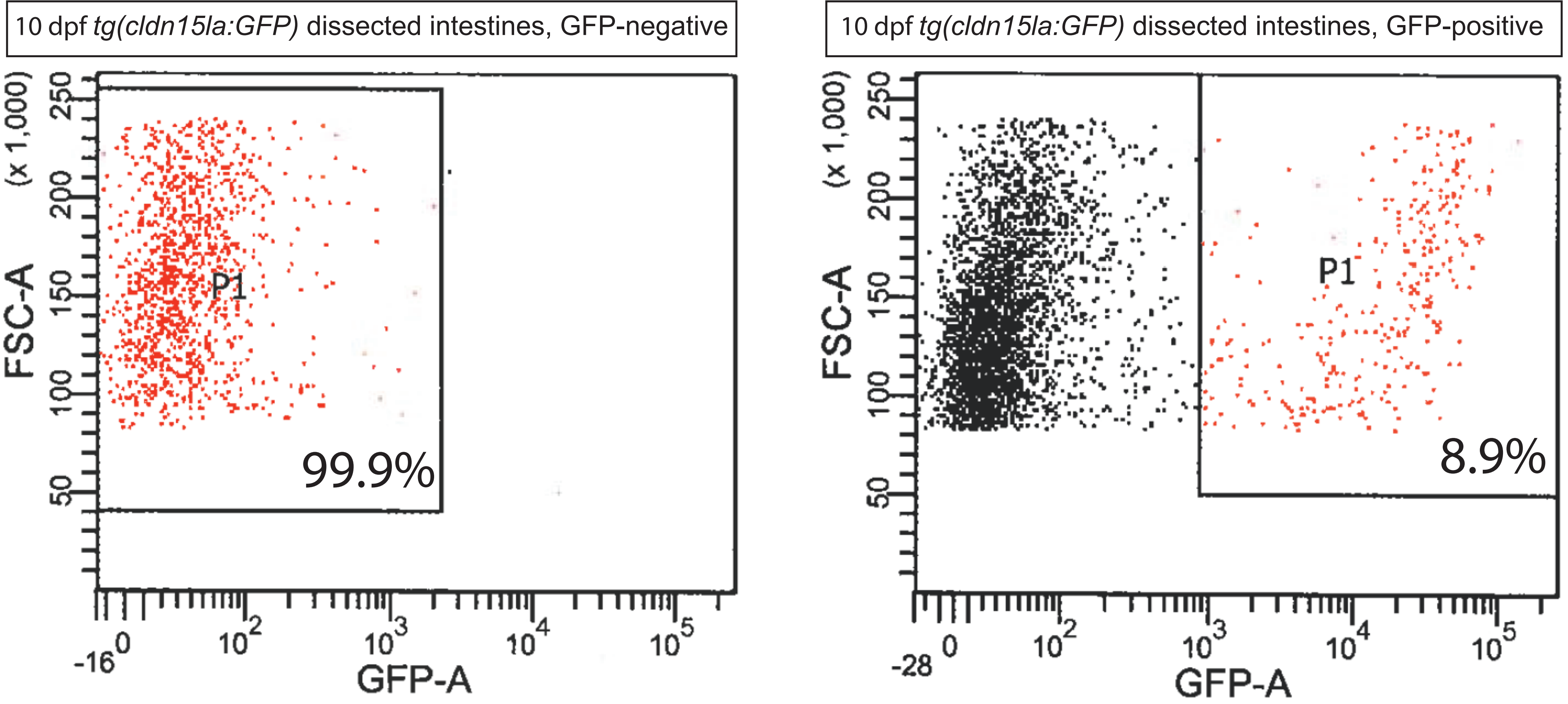


Supplemental Figure 1
